# A heterochromatic histone methyltransferase lowers nucleosome occupancy at euchromatic promoters

**DOI:** 10.1101/429191

**Authors:** H.M. Chen, T.B. Sackton, B. Mutlu, J. Wang, S. Keppler-Ross, E. Levine, T. Liu, S.E. Mango

## Abstract

H3K9me3 (histone H3 modified with tri-methylation at lysine 9) is a hallmark of transcriptional silencing and heterochromatin. However, its global effects on the genome, including euchromatin, are less well understood. Here we develop Formaldehyde-Assisted Identification of Regulatory Elements (FAIRE) for *C. elegans* to examine the chromatin configuration of mutants that lack virtually all H3K9me3, while leaving H3K9me1 and H3K9me2 intact. We find that nucleosomes are mildly disrupted, and levels of H3K9me2 and H3K27me3 rise in mutant embryos. In addition to these expected changes, the most dramatic change occurs in euchromatin: many regions encompassing transcription start sites (TSSs) gain an average of two nucleosomes in mutants. The affected regions normally lack H3K9me3, revealing a locus non-autonomous role for H3K9me3. Affected TSSs are associated with genes that are active in epithelia and muscles, and implicated in development, locomotion, morphogenesis and transcription. Mutant embryos develop normally under ideal laboratory conditions but die when challenged, with defects in morphogenesis and development. Our findings reveal that H3K9me3 protects transcription start sites within euchromatin from nucleosome deposition. These results may be relevant to mammals, where diseases that disrupt the nuclear lamina and heterochromatin can alter epithelial and muscle gene expression.

## Introduction

Chromatin in eukaryotes can be broadly divided into transcriptionally competent euchromatin and less transcriptionally active heterochromatin. A defining feature of heterochromatin is methylation of histone H3 lysine 9 (H3K9me), which promotes chromatin compaction and phase separation in conjunction with H3K9me3 binding proteins (Politz et al. 2013; Strom et al. 2017; Larson et al. 2017). Although genes can transition between heterochromatin and euchromatin, these two states are often viewed as functionally separate entities. Here we test this assumption by asking if disruption of H3K9me3 affects the genome beyond its heterochromatin domains.

H3K9 can be methylated to carry a single, double or triple methyl-group (me1, me2, me3). Fission yeast has a single enzyme that generates all three forms, and is responsible for heterochromatin nucleation, spreading and maintenance (Zhang et al. 2008). As genome complexity increases in higher eukaryotic organisms, H3K9 methylation plays more diverse roles, modifies a larger portion of the genome, and relies on more enzymes for different forms of H3K9 methylation (Allis and Jenuwein 2016). The abundance of H3K9 methylating enzymes in metazoans has made it difficult to understand the function of each methylation state *in vivo*. A recent study using *Drosophila* mutated lysine 9 to arginine, to block all methylation and acetylation (Penke et al. 2016). Nucleosomes were disrupted and expression of piRNAs and transposons increased, however it was impossible to know which modification was responsible for which function. An independent study found that H3K9me2 and H3K9me3 were functionally distinct in *S. pombe* where H3K9me2 domains were transcriptionally active and coupled RNAi-mediated silencing to establish silent H3K9me3 domains (Jih et al. 2017). In this study, we examine the defects associated with selective loss of H3K9me3, leaving H3K9me1 and H3K9me2 intact.

Two key enzymes control H3K9 methylation during *C. elegans* embryogenesis: *met-2* is required for H3K9me1 and H3K9me2, and *set-25* is required for H3K9me3 (Andersen and Horvitz 2007; Towbin et al. 2012; McMurchy et al. 2017). This division of labor enabled us to examine the function of H3K9me3 without eliminating H3K9me1/2. The prevailing view of H3K9me3 predicted that removal of H3K9me3 by *set-25* mutant would destabilize nucleosomes at regions that are normally modified with H3K9me3, leading to de-repression of heterochromatin loci and repeats (Ahringer and Gasser 2018). In contrast to this prediction, we find that disruptions to heterochromatin are relatively minor. Instead, the most prominent changes occur in euchromatin, within cell-type specific promoters, particularly those of epithelial and muscle genes. A large proportion of these promoters gain nucleosomes in mutant embryos, their transcription is disturbed and survival is less robust when environmentally challenged. Our observations suggest that H3K9me3 has dual roles: a direct role in gene silencing and heterochromatin, and a non-autonomous role in gene activation within euchromatic compartments.

## Results

### FAIRE for *C. elegans*

To track the nucleosome configuration in wild-type vs. *set-25* mutants, which lack H3K9me3 (Ashe et al. 2012; Towbin et al. 2012), we developed FAIRE for *C. elegans*. FAIRE uses phenol-chloroform extraction after chemical crosslinking and physical shearing, to isolate DNA that lacks stably associated nucleosomes (Figure 1A; (Giresi and Lieb 2009)). FAIRE is a sensitive indicator of transcriptional activity, by detecting nucleosome-depleted DNA at transcription start sites (TSSs). FAIRE also reveals nucleosomal coverage throughout the genome: low FAIRE reads reflect nucleosome occupancy while high FAIRE reads reflect regions with fewer nucleosomes and/or unstable nucleosomes (Simon et al. 2012; Hogan et al. 2006; Giresi et al. 2007). FAIRE has two advantages compared to other methods such as ATAC or DNase digestion: first, it does not require isolation of nuclei away from the cytoplasm, which could perturb chromatin. Second, it does not rely on enzymatic activities to fractionate DNA, thereby allowing a more linear, quantitative read out; MNase digestion, for example, can produce different results depending on digestion conditions, due to variable nucleosome sensitivity (Mieczkowski et al. 2016; Jeffers and Lieb 2016)(our unpublished observations). FAIRE facilitated comparisons between embryos with different genotypes and different nucleosome configurations. In addition, we included a spike-in control of *C*. *briggsae* embryos and separated *C. elegans* and *C. briggsae* reads computationally (Figure 1A). Mixing species enabled better quantitation between different samples and genotypes (see Materials and Methods).

**Figure 1.**
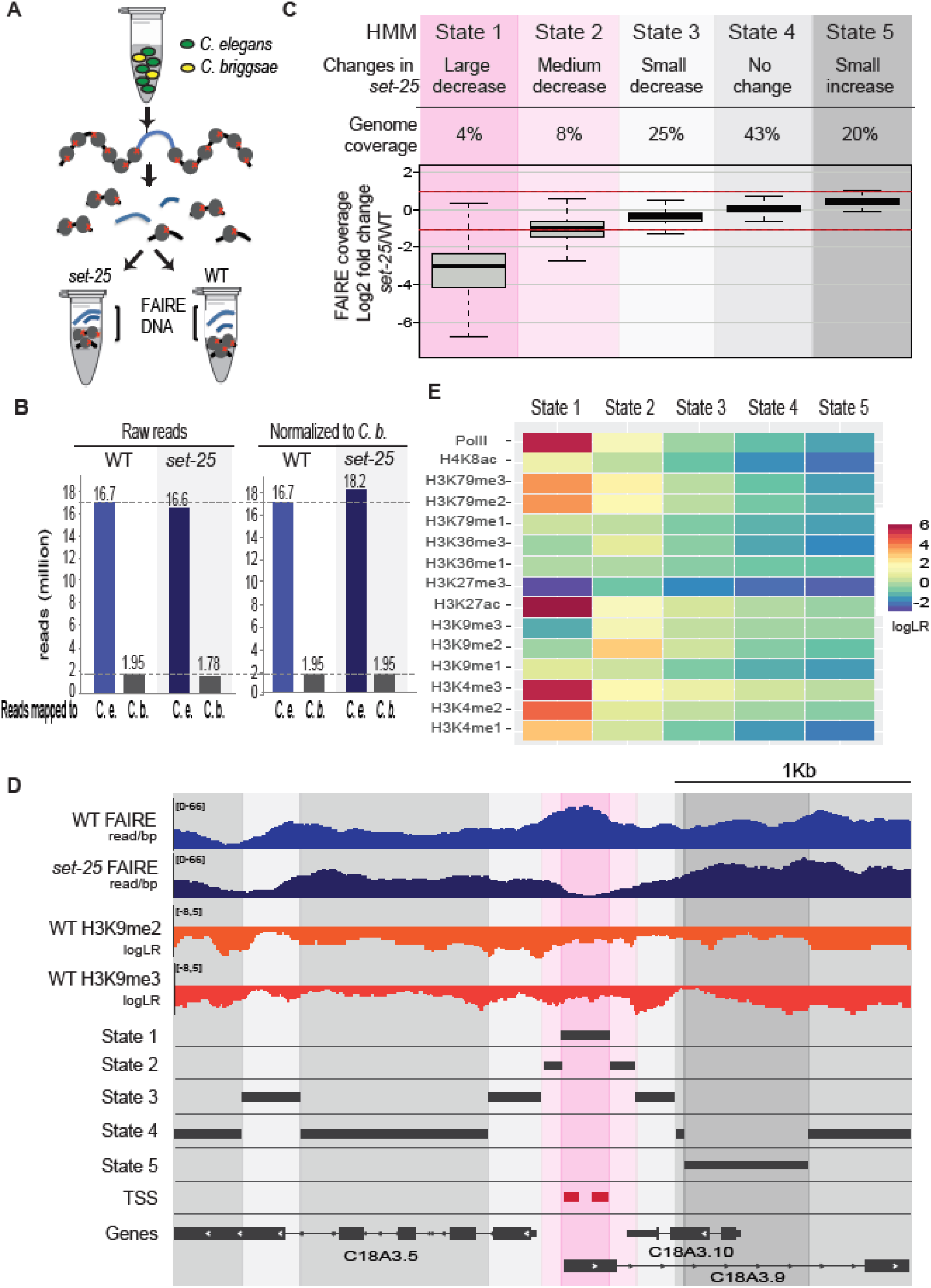
FAIRE analysis identifies nucleosomes sensitive to SET-25 activity within euchromatin. (A) Quantitative FAIRE. Mixed-stage *C. briggsae* embryos (yellow) were distributed among the samples of *C. elegans* embryos (green) at a 1:10 ratio, calculated from the concentration of DNA. FAIRE was conducted on the mixed preparations, and species were sorted computationally after sequencing. This approach normalized for chromatin preparation and sequencing. (B) *C. briggsae* normalization to quantify wild-type (WT) vs. *set-25* sequencing data. (C) Five-state Hidden Markov Modeling (HMM). The genome was categorized into five HMM states based on the different FAIRE values between wild-type and *set-25*. The FAIRE values in log_2_ (*set-25* over wild-type) of all regions in each state are plotted, where the box encompasses the first to third quantiles, the midline reflects the median, and the whisker represents ±1.5x of the Inter Quantile Range. The largest effect was a decrease in FAIRE values for State 1. (D) Heatmap showing the enrichment patterns of RNA polymerase II and 15 histone modifications (Ho et al., 2014, Hsu et al., 2015) over the five HMM states. Note that State 1 is enriched for active marks. (E) A genomic region that contains all five HMM states. X axis: location in the genome. Tracks from top to bottom: Normalized FAIRE reads per base-pair (bp) for i) WT and ii) *set-25* mutant embryos where the Y axis represents the number of FAIRE reads per base pair; note that State 1 has high FAIRE in the wild type and low in *set-25*. Log likelihood ratio (logLR) transformed WT iii) H3K9me2 and iv) H3K9me3 ChIP-seq (Ho et al., 2014) where accumulation below the midline represents depletion and above represents enrichment; note that this region is depleted of H3K9me. v) State 1 to 5 HMM regions denoted by black bars, with State 1 marked in pink. vi) High confidence transcription start sites (red) from (Kruesi et al. 2013) and genes. Gene structure from version ce10.

We performed three biological replicates of FAIRE and focused on regions that were reproducibly different. As a control, we compared the FAIRE output to published MNase studies for nucleosome positioning (Ercan et al. 2011) and found that FAIRE signals reflected the inverse of MNase reads, as expected (Figure S1). On the other hand, our FAIRE counts were only moderately anti-correlated with predicted nucleosome occupancy values, derived from Eran Segal and colleagues (Kaplan et al. 2009). The R value ranged from −0.44 to −0.36 for window sizes spanning 150bp to 5kb, respectively (Figure S1). These data demonstrate that FAIRE worked as expected to define regions of the genome lacking stable nucleosomes, but that nucleosome-depleted regions *in vivo* do not conform closely to computational predictions.

### The *set-25* methyltransferase is required for open chromatin in euchromatin

H3K9me3 promotes nucleosome stability *in vitro* and provides a high-affinity binding motif for heterochromatin protein 1 (HP1) (Zentner and Henikoff 2013; Canzio et al. 2013; Garrigues et al. 2014). To determine if H3K9me3 stabilizes nucleosomes *in vivo*, we generated FAIRE samples from wild-type and *set-25* mutant embryos at equivalent developmental stages (550 cells, bean-1.5 fold, late stage); *set-25* lacks detectable H3K9me3 in embryo samples while retaining H3K9me1 and H3K9me2 ((Towbin et al. 2012); see below). Careful quantitation using the *C. briggsae* spike in revealed that *set-25* late-stage embryos had 9% more FAIRE reads, suggesting a small increase in nucleosome-free DNA compared to wild-type embryos (e.g. 16.7 wild-type vs 18.2 *set-25* normalized million reads; Figure 1B).

To identify regions harboring altered nucleosomal coverage, sequenced FAIRE fragments were analyzed by Hidden Markov modeling (HMM) (Methods). The HMM approach differed from standard analytical methods by identifying regions quantitatively rather than by peak calling, and by allowing the algorithm to define window sizes by tracking FAIRE values at contiguous bases rather than relying on fixed window sizes. This approach provided a sensitive means to compare the entire genome between wild-type and mutant animals.

The HMM analysis categorized the genome into five states (Figure 1C). A genomic region that contained all five states is shown in Figure 1D. The most dramatic difference between wild-type and *set-25* embryos was observed for State 1, which comprised regions that were open, or had high FAIRE values, in the wild type and were nucleosome-occupied in the mutant. State 1 covered 4% of the genome and mapped to euchromatin, regions that were depleted of silencing histone marks and enriched for marks of active transcription and RNA Pol II (Figure 1E). State 2 regions had a similar trend to State 1, but a smaller magnitude, and many State 2 regions (77.5%) were located adjacent to State 1 regions. State 3 had a small decrease in FAIRE values; State 4 had no change between wild-type and mutant. State 5 had more FAIRE reads in mutants compared to wild-type, indicating a slight destabilization or loss of nucleosomes over a large portion of the genome (20%). State 5 presumably accounts for much of the small, overall increase in reads for *set-25* (Figure 1B). Interestingly, State 5 regions were not enriched with H3K9me3 or other silencing marks in the wild type, rather State 5 regions were low for every histone modification, similar to the “Low signal states” identified across metazoan genomes including that of *C. elegans* (Ho et al. 2014).

In sum, the regions affected most dramatically by loss of H3K9me3 mapped to active euchromatin and lead to the addition of nucleosomes. We did not find evidence that depletion of H3K9me3 generated high FAIRE reads, i.e. dramatic nucleosome loss or destabilization, but only a minor disruption, even over repeat regions (described below). This result reveals that H3K9me3 is not a strong determinant for nucleosome stability in *C. elegans* embryos.

### State 1: *set-25* is required for nucleosome-depleted regions at transcription start sites

Given its large effects, we focused on State 1 and characterized State 1 regions in three ways. First, we asked where these regions were located relative to annotated genes (Figure 2A). 92% were positioned within 3 Kb upstream of the nearest gene, and 68% were within 1Kb of the first ATG, including promoter sequences and transcription start sites (TSSs). This bias suggested a possible relationship between State 1 and cis-regulatory sites. Second, we analyzed the size of the State 1 regions (Figure 2B). A nucleosome wraps around 147bp of genomic DNA. 95% of these regions were within 150bp to 1000bp, with an average of 371bp and mode of 271bp. The size suggests that State 1 regions covered an average of two nucleosomes, with a range from one to six. Third, we assigned each State 1 region to the closest gene or to the gene designated by TSS mapping studies (Chen et al. 2013; Kruesi et al. 2013). This approach returned 4,404 unique genes. Figure 2C shows the top 5 Gene Ontology (GO) terms for these genes, ranked by significance. The first and second GO terms were Larval Development and Late Embryo Development, suggesting that *set-25* affects nucleosomes for genes important in development. The third GO term was Locomotion, including cell migration and organism movement. The fourth category was Body Morphogenesis, a process that relies heavily on forces generated by epithelia and muscles to elongate the embryo from a ball of cells into a tube-shaped worm in late-stage embryos (Priess and Hirsh 1986; Vuong-Brender et al. 2016). The fifth term was Positive Regulation of Transcription. Together, these GO terms suggest that *set-25* affected genes active in late embryogenesis. The stage parallels the stage when embryos were harvested for FAIRE, suggesting active *set-25* affected active genes.

**Figure 2.**
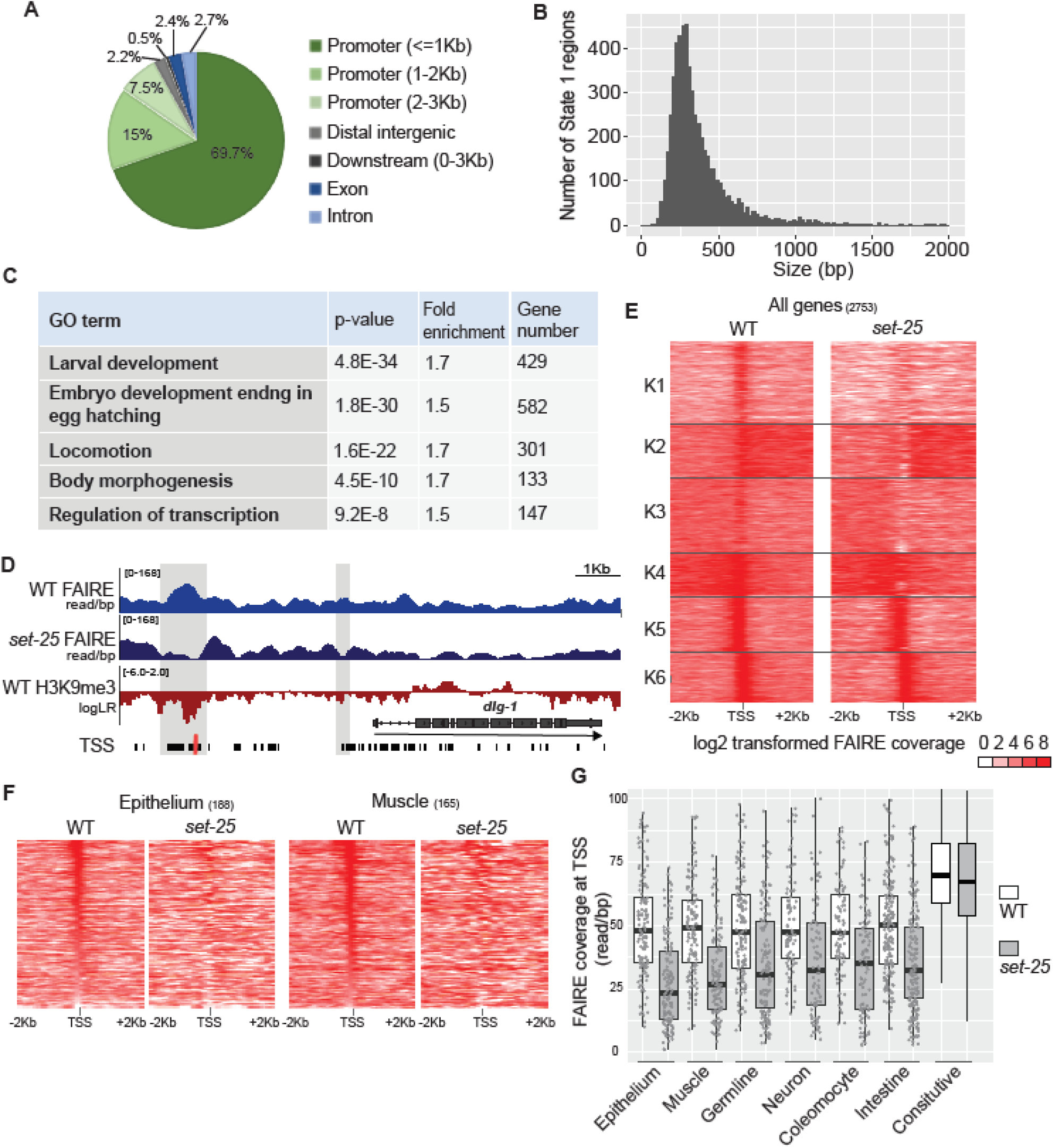
*set-25* is required for nucleosome-free regions at transcription start sites. (A) Genome distribution of State 1 HMM regions, which are enriched for Promoter regions. (B) Size distribution of State 1 regions. (C) GO term analysis for genes classes associated with State 1. (D) Example of a State 1 gene with reduced FAIRE signals in *set-25* mutants. Upper tracks (blue): WT and *set-25* normalized FAIRE values. Lower track (red) LogLR transformed WT H3K9me3 ChIP-seq from Ho et al., 2014. State 1 regions are shaded in grey. Two TSS tracks are shown below the *dlg-1* gene structure (black from Chen et al. 2013, red from Kruesi et al. 2013). (E) Heatmap showing FAIRE coverage centered at the TSS and spanning 4 Kb. The log_2_ FAIRE coverage spans from zero (white) to 8 (red), and each horizontal line represents one gene. TSSs listed here were identified by both Chen et al., 2013 and Kruesi et al., 2013. Genes were grouped by K-means clustering. (F) Heatmaps for genes enriched in epithelia (left) or muscle (right) for WT and *set-25* embryos, spanning 300bp and centered at the TSS. Genes are sorted by their WT FAIRE values, and the same order is used to plot *set-25* values. Gene lists are derived from (Spencer, 2011, Genome Research). Note the loss of FAIRE values in *set-25* mutants, indicating acquisition or stabilization of a nucleosome over the TSS. (G) Box plots of FAIRE values for 300 bp around the TSSs of cell-type enriched genes. Gene lists are derived from (Spencer, 2011, Genome Research). Genes in the “Constitutive” category are those in clusters K5 and K6 of Figure 2E. Each dot represents the FAIRE value at 100bp around the TSS for each gene.

An example of a State 1 gene is *dlg-1/discs large*, a junctional protein expressed in epithelia (Figure 2D) (Bossinger et al. 2001; Firestein and Rongo 2001; McMahon et al. 2001). *dlg-1* is a member of three of the GO terms associated with State 1: Larval Development, Late Embryo Development and Body Morphogenesis. Two State 1 HMM regions were assigned to *dlg-1*, and each of these overlapped with previously defined TSSs and promoter regions (Figure 2D). To determine the generality of this result, we compared State 1 regions to two embryonic TSS datasets, one generated by Chen et al., and one by Kruesi et al. (Chen et al. 2013; Kruesi et al. 2013). The Chen dataset is comprehensive, with many TSSs (e.g. Figure 2D). The Kruesi dataset is more restricted, with one high-confidence TSS assigned per gene (e.g. Figure 2D). We generated a combined TSS list for 2753 genes with TSSs that had been identified in both the Chen and Kruesi datasets, and we plotted the FAIRE signal over 4kb, centered at the TSS (Figure 2E). K-means clustering revealed six categories of TSS behavior. FAIRE signals for K1, K2 and K3 had medium FAIRE levels in the wild type, suggesting that normally, these TSSs were open in a subset of cells or open in many cells but with low frequency; most of these TSSs lost FAIRE signals in *set-25* mutant embryos. On the other hand, K5 and K6 had high FAIRE values for wild-type TSSs, and these were largely unaffected by *set-25* mutations (Figure 2F, G). We envision that these genes are likely active in many cells or expressed strongly. This result suggests that *set-25* had greater impact on cell-type specific or weakly expressed genes.

To determine whether *set-25*-sensitive genes reflected genes expressed in a particular cell type, we examined lists of genes expressed in epithelia, muscles, germ line, neurons, coelomocytes and intestine (Spencer et al. 2011). Genes in each of these sets were strongly affected by *set-25*, particularly those in the epithelium and muscle groups, compared to the constitutive FAIRE classes (Figure 2G). We conclude that *set-25* is required to establish nucleosome-depleted regions at the TSSs of cell-type specific genes.

### *set-25* mutants have reduced RNA expression

Generally, TSSs depleted of nucleosomes reflect active or poised promoters, and many of the regions affected by *set-25* mutants are normally active in mid to late-stage embryos (Adelman and Lis 2012). The loss of FAIRE coverage at TSSs in *set-25* mutants suggested that nucleosomes might hinder transcription for these loci. To test this idea, we examined RNA expression for three epithelium genes that acquired closed TSSs in *set-25* mutants (*dlg-1, chtl-1* and *sqv-3*), one housekeeping gene with an open TSS in both wild-type and *set-25* (*eft-3*), and one gene marked by H3K9me3 in the wild type *etr-1* (Figure 3B). To obtain the most quantitative measurements of mRNA abundance, we used digital droplet PCR on single animals (Figure 3A; (Sykes et al. 1992)). We analyzed both late-stage embryos, at the time embryos were harvested for FAIRE, and first-stage larvae after hatching (Figure 3C and D).

**Figure 3.**
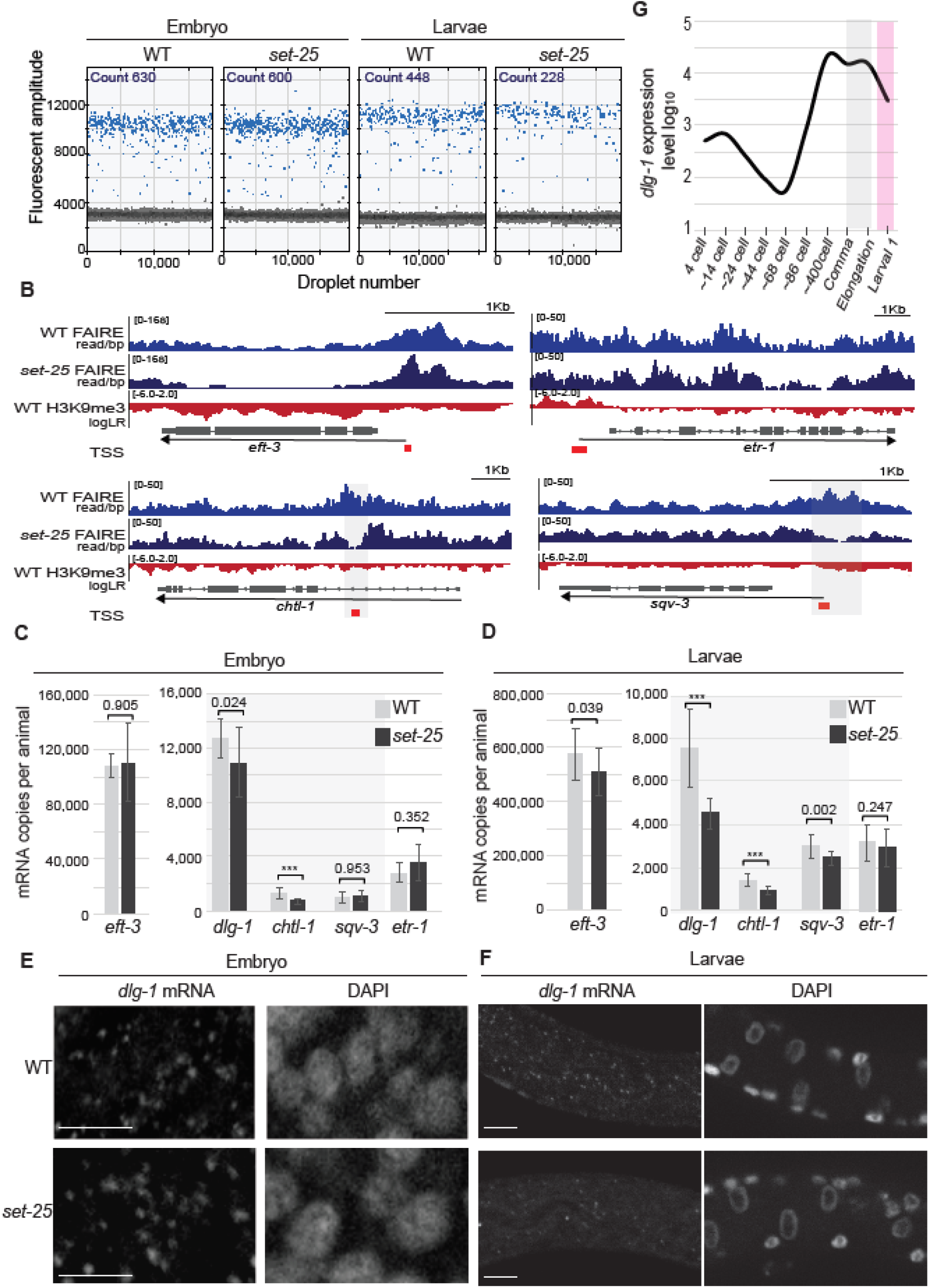
Decreased mRNA levels for epithelium genes in larvae. (A) Digital droplet PCR counts for *dlg-1*. Each cDNA sample was partitioned into ∼20,000 PCR reaction droplets. The blue dots had high fluorescent amplitude and were considered positive. The black dots were considered negative. The number of positive droplets per sample is noted (count).(B) Genome regions for *eft-3* (control)*, chtl-1, etr-1*, and *sqv-3*. Upper tracks (blue): WT and *set-25* normalized FAIRE values from embryos. Lower track (red) LogLR transformed WT H3K9me3 ChIP-seq. Altered State 1 FAIRE regions are shaded grey for *chtl-1 and sqv-3*. The gene structure is based on ce10 and the TSS denoted in red. (C) Digital droplet PCR counts for 16 individual embryos (1.5 fold). Statistical t-test p-value is labeled for each wild type and *set-25* reaction pair. *** indicates p value < 0.005.(D) Digital droplet PCR for 16 individual L1 larva, quantified as in C. (E) Single molecule FISH (smFISH) for *dlg-1* mRNA in embryos. Left: each dot represents a single *dlg-1* mRNA. Right: DAPI stain of nuclei. These are foregut cells. Scale bar = 2 um.(F) smFISH for *dlg-1* mRNA in first-stage larvae, marked as in E. These are midgut cells. Scale bar = 2 um.(G) Quantitative microarray data for *dlg-1* mRNA adapted from (Levin et al. 2012). Grey box: time when TSSs lost FAIRE signal in *set-25* mutant embryos. Pink box: time when decreased expression of epithelium genes was observed.

In embryos, we observed only minor changes in *set-25* mutants, and these were not statistically significant (Figure 3C, E). However, multiple genes were affected in L1 larvae (Figure 3D, F). The three epithelial factors *dlg-1, chtl-1* and *sqv-3* all had statistically significant decreases in RNA expression, ranging from 20% to 40%. The heterochromatin gene *etr-1* remained unchanged in mutants compared to wild-type, consistent with the FAIRE data (Figure 3B). *eft-3* had a small decrease that was not statistically significant; the *eft-3* decrease did not correlate with decreased expression of epithelial genes on a single-worm basis (Figure 3C, D). We interpret this result to mean that individual animals had variable expression of *eft-3* and other genes, and not that lower values of *eft-3* reflect a lower yield of RNA in a given animal. These data suggest that epithelial gene expression is selectively reduced in *set-25* mutants and variable from animal to animal. To test this idea further, we used single-molecule fluorescent *in situ* hybridization, smFISH, to visualize *dlg-1* mRNA in individual cells at specific stages. Consistent with the results from digital droplet PCR, *dlg-1* mRNA from *set-25* mutants was similar to wild type in embryos but reduced in larvae (Figure 3E, F). These findings suggest that aberrant nucleosome deposition in *set-25* mutants leads to reduced expression of epithelial genes, but that there is a delay between TSS occupancy and RNA accumulation.

#### *set-25* mutants are sensitive to hypo-osmotic pressure

To date, no morphogenesis defect has been described for *set-25*, but we speculated that decreased epithelial gene expression could affect the biology of *set-25* mutant worms. Under ideal laboratory conditions, *set-25* mutants grow like wild-type: 93% of *set-25* mutants completed embryogenesis, compared to 96% for the wild type (Figure 4A); *set-25* mutant embryos developed from the 2-cell stage to the L1 stage in approximately 750 ±14 minutes at 20°, compared to 740 ±17 minutes for the wild type (Figure 4B); *set-25* adult worms had the same brood size as wild-type embryos (Figure 4C). None of these differences between *set-25* and wild-type were statistically significant, suggesting that development is buffered against small changes in gene expression for *set-25* mutants grown in ideal conditions.

**Figure 4.**
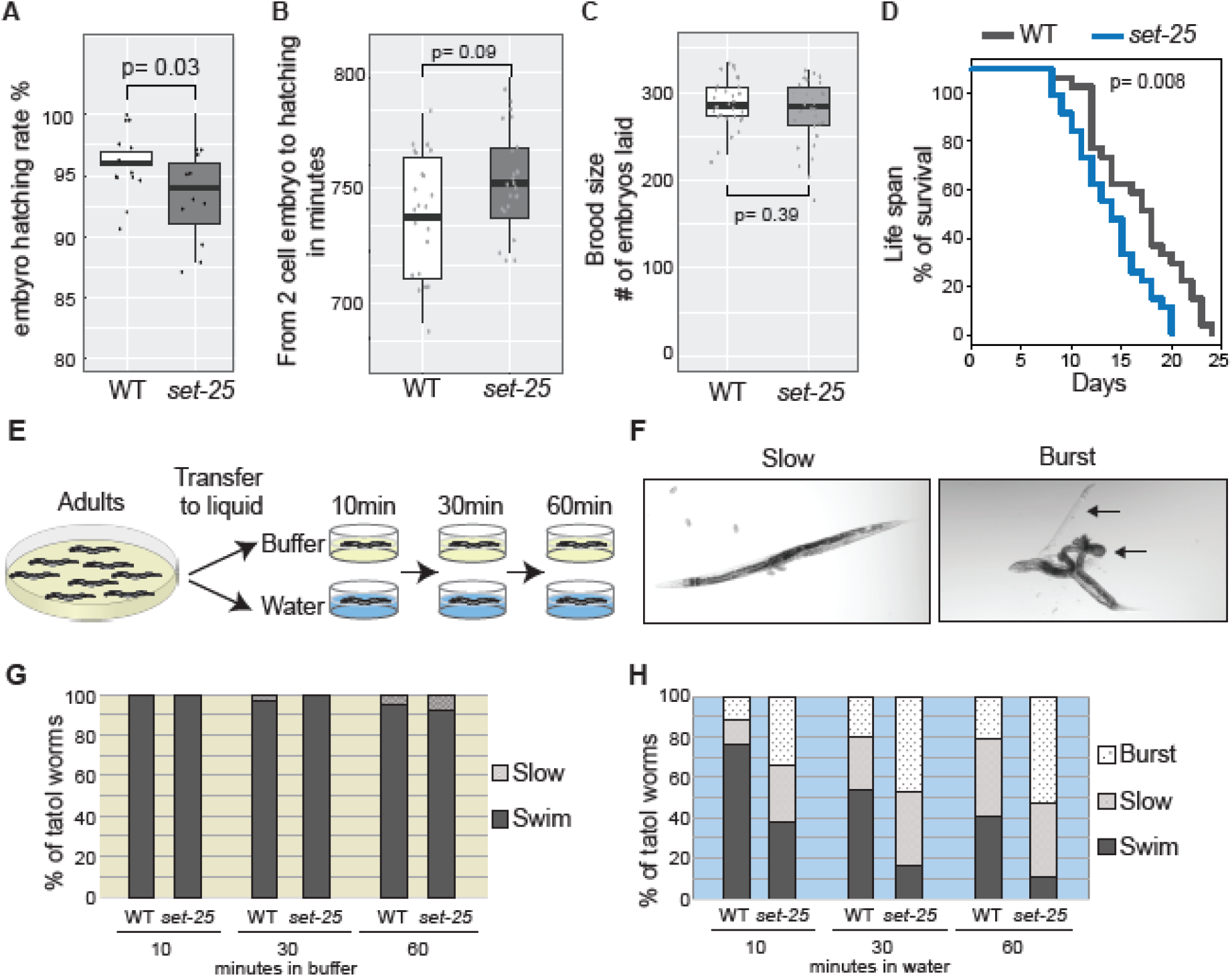
*set-25* mutants are sensitive to hypo-osmolarity. (A) Embryo hatching. Each dot resents an individual experiment of 25 embryos. 4 independent experiments per genotype. p-value of t-test is shown. *set-25* mutants were largely viable. (B) Time from 2-cell embryo to hatched L1 larva. Each dot represents an individual animal. 10 animals for each genotype per experiment, and two experiments. (C) Brood size. Each dot represents the offspring from one individual animal. 10 animals per genotype per experiment and three experiments. (D) Life span. Each line represents the combined results of 30 animals in three experiments. 10 animals per genotype per experiment, and three experiments. Note that *set-25* had decreased longevity compared to the wild type. (E) Rainy day challenge. Adult worms were transferred to M9 salt buffer (no osmotic pressure) or to water (hypo-osmotic pressure), and scored after 10, 30, or 60 minutes. (F) Bright field image of *set-25* adults in water. Slow: Worms were either visibly slower than normal, or were paralyzed completely. These worms maintained a worm structure. Burst: The gonad (upper arrow) and the midgut (lower arrow) erupted through the vulva. (G) Phenotype in M9 salt buffer. Bars represent the combined results of three experiments with 48 worms per experiment, in triplicate. Almost all worms survived. (H) Phenotype in water. Bars represent the combined results of three experiments with 48 worms per experiment, in triplicate. Note that set-25 was considerably more sensitive to incubation in water.

We observed one difference in the unperturbed life cycle of wild-type vs. *set-25* worms. Adult *set-25* mutants lived approximately three days shorter than wild-type animals (Figure 4D). This result is intriguing, given that H3K9me3 is reduced during normal aging (reviewed in (Benayoun et al. 2015)). The effect of *set-25* extends the list of histone methyltransferases that modulate life span (Benayoun et al. 2015).

When challenged by less ideal growth conditions, *set-25* mutants exhibited less resilience compared to wild-type worms.*C. elegans* grow in damp environments in the wild (i.e. soil, fruit). The standard laboratory strain Bristol was isolated from Bristol, UK, where it rains an average 18.6 days per month (weather.com). We replicated the rainy-day environment by transferring age-matched adults to water for 10, 30 or 60 minutes (Figure 4E). Wild-type animals survived at least two-fold better compared to *set-25* animals in this environment. Whereas 75% of wild-type animals swam vigorously when incubated for 10 minutes in water, less than 40% of *set-25* mutants swam normally. The remainder either became rigid and stopped swimming, or they burst, with the midgut and gonad erupting through the vulva (Figure 4G, H). At 60 minutes, the phenotype was even more pronounced, with only 10% of *set-25* mutants able to swim normally, compared to 40% of wild-type worms. The bursting phenotype suggested that the cuticle and epidermis were less robust in *set-25* mutants and succumbed to the build-up in osmotic pressure. Consistent with this hypothesis, both wild-type and *set-25* mutant adults survived well in M9 buffer, which contained 150mM salt, compared to the water assay, which lacked salt altogether (Figure 4G). These data suggest that defects in chromatin and gene expression in *set-25* mutants produce defects in the worms’ protective barrier of the epithelium and cuticle.

### *set-25* is required for robust embryogenesis when challenged

The water assay challenged the external epidermis but did not probe internal epithelia. As a test of internal epithelial formation and integrity, we examined the ability of embryos to withstand ectopic expression of a developmental regulator. We chose the potent transcription factor HLH-1/MyoD (Fukushige and Krause 2005) and focused on late-stage embryos, which had altered FAIRE landscapes in *set-25* mutants. Briefly, *hlh-1* was activated ubiquitously in late-stage embryos using a heat-shock promoter. After recovery, embryos were examined either at an intermediate time point (5 hours) or at the end of embryogenesis (16 hours; Figure 5A).

**Figure 5.**
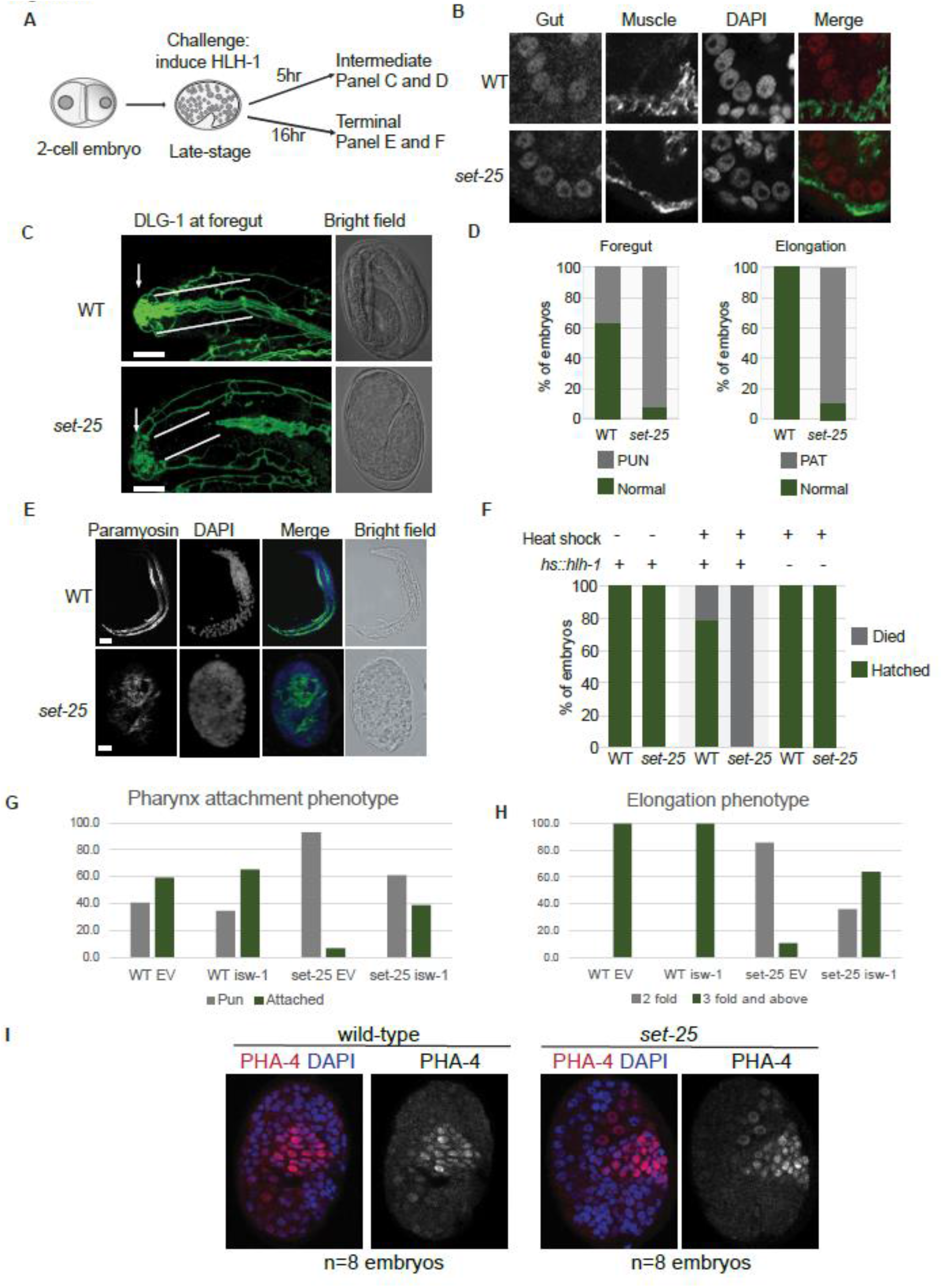
*set-25* is required for robust embryogenesis when challenged. (A) Experimental scheme of HLH-1 challenge. Wild-type or *set-25* embryos carried *hlh-1* under control of the heat-shock promoter. 2-cell stage embryos were collected, aged (to the comma stage), and heat shocked at 34°C for 10min to induce HLH-1 ectopically. Embryos were allowed to recover for 5 or 16 hours before examination. (B) Wild-type (upper) and *set-25* mutants (lower) express gut (PHA-4, red) and muscle (paramyosin, green) markers appropriately. DAPI marks DNA. (C) 5 hour incubation, intermediate phenotype: left panels: embryos stained for epithelia (DLG-1; green). Wild-type embryos have a foregut epithelium attached to the buccal cavity (arrow; (white lines) whereas *set-25* mutant do not (note the lack of green between the white lines; pharynx unattached or Pun phenotype). Right panels: light microscopy images. WT embryo elongated past the 3-fold, but the *set-25* mutant arrested at the two-fold. Scale bar = 5m. (D) Quantitation of the foregut and elongation phenotypes. PUN= foregut unattached. PAT= twofold arrest. N= 3 experiments, 27 embryos for WT, 28 embryos for *set-25*. Note the *set-25* has more PUN and more PAT embryos. (E) 16 hour incubation, terminal phenotype: embryos were stained for muscle (paramyosin, green), DNA (DAPI, blue). Light microscopy shows the lack of elongation for *set-25* (lower) compared to te wild type (upper). (F) The rate of hatching for unchallenged (- heat shock) and challenged (+ heat shock) embryos. Embryos carried the hs::hlh-1 transgene (+) or not (-). Note that *set-25* mutants were highly sensitive to *hs::hlh-1* induction (middle columns) compared to wild-type or controls. Died= embryos lost their normal morphology and arrested. N= three experiments. Embryos number for each column, left to right: 56, 57, 28, 33, 23 and 25. (G) Rescue of PUN animals after hs::hlh-1 by isw-1(RNAi) compared to RNAi control (EV). Y axis: % embryos with a Pun foregut (grey) or a normal, attached foregut (green). Note that *set-25* mutants are normally Pun after treatment, but are rescued when the chromatin remodeler *isw-1* is inactivated. (H) Rescue of body elongation after hs::hlh-1 by isw-1(RNAi) compared to RNAi control (EV). Y axis: % embryos that elongated fully (green) or arrested at the 2-fold (grey). Note that most *set-25* mutants arrest after treatment, but are rescued when the chromatin remodeler *isw-1* is inactivated. (I) Foregut specification occurs normally in wildtype (left) or set-25 (right) embryos after HS::hlh-1 treatment. Foregut cells stained for PHA-4 (pink).

Embryo morphogenesis was disproportionately disturbed in *set-25* mutants. At the intermediate time point, challenged *set-25* mutant embryos were defective for body elongation, a process in which nascent epithelia and muscles mold ball-shaped embryos into tube-shaped worms (3-fold stage;(Sulston et al. 1983; Priess and Thomson 1987). All wild-type embryos elongated beyond the 2-fold stage (Figure 5C, D). However, only 10% of *set-25* mutant embryos elongated properly, with the remaining 90% resembled a 2-fold stage embryo (Figure 5C and 5D). 16 hours after HLH-1 induction, the wild-type embryos elongated fully and 80% of them hatched into L1 larvae (Figure 5D and 5E). No *set-25* embryos hatched after HLH-1 induction, and many reverted to a clump of cells through failed morphogenesis. We observed a second morphogenesis defect in the gut. 63% of wild-type embryos formed a linear digestive tract, with the foregut attached to the buccal cavity, whereas only 7% of *set-25* mutants made a linear gut. The remainder were Pun (pharynx unattached), with an absence of epithelial markers in the anterior, which indicated that the tip of the foregut was not attached to the buccal cavity (Figure 5C and 5D).

Next, we examined survival after HLH-1 induction in combination with inactivation of *isw-1/imitation switch*. *Imitation switch* is a nucleosome remodeler that moves nucleosomes at TSSs and can promote the reformation of chromatin (Adelman and Lis 2012). We reasoned that if Histone H3 is the relevant substrate of the *set-25* enzyme, we should observe genetic interactions between *set-25* embryos and *isw-1* after heat shock. Indeed, *set-25* mutants survived better when *isw-1* was inactivated by RNAi compared to a control RNAi (Figure 5G, H). Wild-type embryos were not affected by *isw-1(RNAi*), suggesting that the effect was specific (Figure 5G, H).

We performed two follow-up controls for these experiments. First, to ensure that the poor survival of *set-25* mutants reflected the activity of the HLH-1 transcription factor, we subjected wild-type and mutant embryos to heat shock in the absence of *hlh-1*. Both wild-type and mutant embryos survived, indicating that the lethal *set-25* phenotype reflected a response to HLH-1 and not to heat shock alone (Figure 5F). As a second control, we stained heat-shocked embryos for the foregut factor PHA-4. Previous studies had shown that PHA-4 is a sensitive read-out for cell-fate conversion (Kiefer et al. 2007; Mango 2009). If HLH-1 converted cells to a body wall muscle identity, we predicted we would see reduced PHA-4 staining. If HLH-1 generated morphological mayhem without cell-fate conversion, we expected that embryos would continue to express PHA-4 normally, even if they failed to undergo morphogenesis. In both wild-type and *set-25* mutants, we observed many PHA-4 positive cells that resembled foregut (high PHA-4 levels; small cells) or midgut (low PHA-4 levels, large cells), suggesting that the increased lethality of *set-25* mutants did not reflect increased cell-fate conversions (Figure 5B, I). Body wall muscles were also appropriately established, suggesting cell fates were established correctly (Figure 5E). These data reveal that challenging cells with ectopic HLH-1 does not alter cell fate, in agreement with previous studies but produces defects in embryonic morphogenesis (Horner et al. 1998; Djabrayan et al. 2012; Zhu et al. 1998; Gilleard and McGhee 2001; Fukushige and Krause 2005; Kiefer et al. 2007; Mango 2009; Priess and Thomson 1987).

### *set-25* is dispensable for heterochromatin silencing

In addition to euchromatin, we examined silent regions of the genome in *set-25* mutants. Like other animals, H3K9me3 is associated with heterochromatin and DNA repeats in *C. elegans* (Gu and Fire 2010; Liu et al. 2011; Gerstein et al. 2010; Garrigues et al. 2014; McMurchy et al. 2017) Previous studies with worms showed that loss of all methylated H3K9 (H3K9me1, H3K9me2, H3K9me3) lead to de-repression of some repeats (Zeller et al. 2016; McMurchy et al. 2017). Similarly, work in *Drosophila* that eliminated all H3K9 modifications with a lysine to arginine substitution found that repeats and transposons were de-condensed (Penke et al. 2016). To determine the contribution of H3K9me3 alone, we examined *set-25* mutants for heterochromatic regions, including repetitive sequences that were modified by H3K9me2/3 and bound by heterochromatin protein HPL-2 (Garrigues et al. 2014). These regions were weakly enriched in State 2.

Surprisingly, most repeats had normal FAIRE values in *set-25* mutant embryos in late-stage embryos. Figure 6A shows a section of the genome with PALTTTAAA3, a non-autonomous DNA transposon that lacks transposase, and CeREP55 mini-satellite repeat clusters. Figure 6B shows a genomic region with the DNA transposon Mirage1 and a Helitron non-autonomous DNA transposon. *set-25* mutations did not alter the FAIRE profile of these repeats and, in particular, there were no TSS peaks to suggest transcriptional induction. These data suggest that repeats remained largely unaffected in *set-25* mutants.

**Figure 6.**
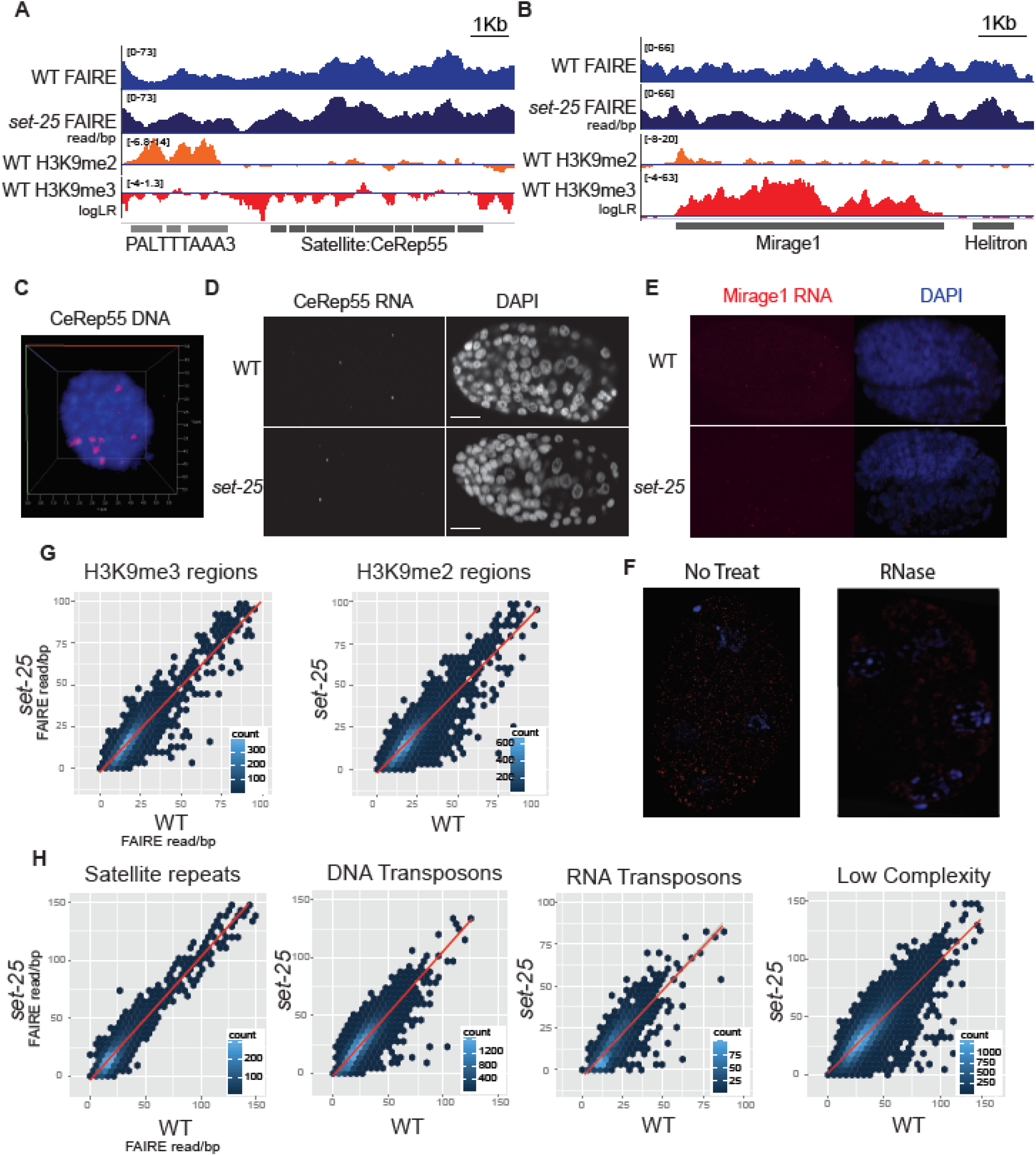
*set-25* has minor impact on heterochromatic regions and repeats. (A) An example region containing PALTTTAAA3 (DNA transposon) and CeRep55 (Satellite repeat). Normalized FAIRE counts per base-pair tracks are shown for WT and *set-25* mutants (blue, upper tracks). LogLR transformed H3K9me2 (orange) and H3K9me3 (red) ChIP-seq tracks i wild-type embryos, where (Ho et al., 2014). Accumulation below the midline represents depletion and above represents enrichment. Note the depletion of H3K9me3 for this region Repeats are marked in grey. (B) An example region containing DNA transposon repeats Mirage1 and HelitronY1. Tracks as in A. Note that Mirage 1 has ample H3K9me3 (C) CeRep55 DNA FISH of single nucleus in 3D. Red spots denote CeRep55 clusters in the genome. X, Y and Z axis are 5m. (D) CeRep55 smFISH of age-matched late stage WT and *set-25* embryos (left) and DNA (DAPI, right). Scale bar= 10m. Note the low abundance of CeRep55 in both strains. (E) Mirage1 smFISH of age-matched WT and *set-25* embryos. Scale bar= 10m. Note the low abundance of Mirage 1 RNA in both strains (F) Mirage1 smFISH of *nrde-2* embryos with and without RNase treatment. Mirage 1 is expressed in *nrde-2* embryos and signal is lost after RNase. (G) High density scatter plot of WT vs. *set-25* FAIRE coverage for regions modified by H3K9me2 (left) and H3K9me3 (right) in the wild type. Each dot represents a collection of regions that fit the FAIRE coverage in WT on the x-axis and *set-25* on the y-axis. The color of the dot represents the count of regions. The red line represents the linear regression. (H) High density scatter plot of WT vs. *set-25* FAIRE coverage for satellite repeats, DNA transposons, RNA transposon and low complexity regions.

As an alternative means of examining repeats, we performed smFISH for Mirage1 and CeRep55 RNA. Mirage1 is heavily decorated with H3K9me3 in the wild type and lacks H3K9me2 (Gerstein et al. 2010). It was a good candidate for derepression by *set-25* mutations. CeRep55 has relatively high FAIRE values in the wild type, but is not enriched for H3K9me2/3 (Gerstein et al. 2010). We imagined that CeRep55 might be de-repressed more easily than other repeat types. Mirage1 and CeRep55 had low expression in *set-25* mutants, and both resembled wild-type embryos (Figure 6D, E). For both repeats, little RNA was detected in nuclei, which is often an indicator of transcription (Batish et al. 2011). We detected more Mirage1 signal in first-stage larvae, particularly in cells that were likely nascent germ-line cells, based on their position (Figure S2). A proportion of both wild-type and *set-25* larvae had germ-line expression. In addition to the bright, punctate signal, we observed a general glow, which may represent degraded or processed RNA (Figure S2). RNAse treatment removed both the puncta and the glow, revealing that our method visualized RNA, as expected (Figure 6F). As an additional control, we examined *nrde-1* mutants, which de-repress Mirage1 in the adult germ line (McMurchy et al. 2017) and detected increased Mirage1 signal (Figure 6F). In sum, these data suggest that loss of H3K9me3 had relatively small effects on Mirage1 and CeRep55 during *C. elegans* embryogenesis or first stage larvae.

To determine if the absence of altered FAIRE values in *set-25* mutants held true for most heterochromatic regions, we plotted FAIRE counts for wild type vs. *set-25* for all H3K9me2 and H3K9me3 marked regions (Figure 6G), and for four repeat classes: satellite repeats, DNA transposons, RNA transposons and regions of low complexity (Figure 6H). Like Mirage1 and CeRep55, most heterochromatic regions had normal FAIRE values in *set-25* mutant embryos, exemplified by the 1-to-1 regression fit line. There were some deviations but many of these appeared to have lower FAIRE values in *set-25* mutants. These data reveal that H3K9me3 is not essential to determine the level of nucleosome coverage in *C. elegans* heterochromatin. This may explain why *set-25* mutants are healthy under ideal laboratory conditions.

A second explanation for the health of *set-25* mutants is the existence of compensatory processes. Consistent with this idea, we observed higher levels of repressive marks H3K9me2 and H3K27me3 in *set-25* mutants (Figure S2). In addition, other silencing pathways may function, such as small RNA mediated silencing, as has been observed in other studies (Gaydos et al. 2014; Martens et al. 2005; McMurchy et al. 2017).

## DISCUSSION

Classical studies of H3K9me3 have focused on its roles in heterochromatin, and silencing genes and repeats (Politz et al. 2013). Here we uncover an additional function, to promote expression within euchromatin. H3K9me3 ensures that active TSSs remain depleted of nucleosomes. In *set-25* mutants, nucleosomes are deposited on TSSs leading to reduced gene expression and lethality upon environmental challenge. Surprisingly, H3K9me3 plays a minor role to configure nucleosomes in heterochromatin. Worms rely on robust silencing pathways, and redundancy between these pathways may ensure that heterochromatic regions can withstand fluctuations or loss of silencing components. Consistent with this idea, loss of H3K9me reduces association of HPL-2/HP1 with chromatin, but does not remove it completely (Garrigues et al. 2014).

The most dramatic change for mutant embryos lacking H3K9me3 was repositioned nucleosomes on the TSSs of cell-type specific genes. Genes that are transcriptionally active, or soon to be active, typically lack a stable nucleosome at their TSS (Adelman and Lis 2012). The correspondence between the age of embryos analyzed by FAIRE and the GO terms for the affected genes suggests that loss of H3K9me3 impacts this cohort of active and primed genes. In *C. elegans*, genes often have many TSSs spread over hundreds of base pairs (McMurchy et al. 2017), and this organization may explain why some regions affected by *set-25* mutations were as large as six nucleosomes, likely spanning many TSSs.

RNA expression and FAIRE were normal in young *set-25* embryos (data not shown), and the reduction of mRNA was only observed after embryo hatching. This result suggests that transcription may initiate normally in *set-25* mutants, but has difficulties with maintenance, growth or reactivation at later stages. A role in reactivation might explain why genes expressed in a subset of tissues were more affected by *set-25*, compared to ubiquitously-expressed genes, which are likely expressed less dynamically. In our RNA survey, *dlg-1* was the most affected by *set-25*, and *dlg-1* has the most dynamic expression profile in wild-type (Figure 3G), compared to ubiquitous genes like *eft-3* or epithelial genes *sqv-3, chtl-1* or *etr-1* (wormbase.org).

An interesting question for future studies is whether reduced H3K9me3 also lowers gene expression in other organisms. It is known that H3K9me3 is dramatically reduced in both Hutchinson-Gilford progeria syndrome (HGPS) and Werner syndrome, due to aberrant nuclear lamins (Kudlow et al. 2007; Prokocimer et al. 2013). Gene expression in skin and muscle are affected in progeria, reminiscent of *set-25* animals, and conversely *set-25* mutants have a shorter lifespan similar to progeria. Unlike lamin mutants, however, the nuclei of *set-25* mutants appear spherical and intact. A speculative but intriguing possibility is that reduction of H3K9me3 explains the defective epithelial and muscle gene expression in these human syndromes.

We also examined repeats and heterochromatin in *set-25* mutants, and found evidence of only minor disruption. FAIRE is a sensitive measure of transcription, and this method complements smFISH and digital droplet RT-PCR. FAIRE can detect transcription from just a few cells ((Giresi et al. 2007); our observations) and is not sensitive to RNA degradation. On the other hand, smFISH and RT-qPCR can detect RNA even after transcription has ceased or from a subset of repeats. By all three measures, we detected repeats and transposons were relatively normal compared to wild-type. Thus, endogenous repeats may be regulated more tightly compared to repeats in extrachromosomal repeat arrays, which are de-repressed by *set-25* ((Towbin et al. 2012); unpublished observations). A likely possibility is that alternative silencing pathways maintain repression in *set-25* mutants. For example, H3K9me1 and H3K9me2 are still present in *set-25* mutants, as is H3K27me3, and levels of these modifications rose in *set-25* mutants. In addition, other SET proteins as well as RNAi pathways can synergize with chromatin regulation to maintain silencing (McMurchy et al. 2017; Ashe et al. 2012). Perhaps defects occur only when these synergistic pathways are disrupted by mutation or by extreme environmental conditions, such as high temperature.

### Materials and Methods

#### Worm maintenance and embryo collection

Worms were maintained on OP50 plates at 20°C for regular husbandry (Brenner 1974). Synchronized embryos were collected from worms grown in liquid culture and incubated at 20°C with shaking at 200rpm in CSM medium without food (100mM NaCl, 5.6mM K_2_HPO_4_, 4.4mM KH_2_PO_4_, 10ug/ml cholesterol, 10mM potassium citrate, 2mM CaCl2, 2mM MgSO4, 1X trace metals). After 16 hours, worms were synchronized to the L1 stage (Stiernagle et. al, 2005). Concentrated NA22 *E. coli* were added and the culture shaken for 62–66hr. Embryos were harvested when ∼50% of the population carried 1–8 embryos, while the rest carried no embryos. These synchronized early embryos were incubated in M9 buffer for 5hr for late-stage with ∼550-cells. Embryos were frozen in liquid nitrogen and stored at −80°C prior to Western blot and FAIRE procedures.

#### Strains

N2 wild type,

MT1746 *set-25(n5021)* III

SM2502 *pxEx627 (let-413::GFP, pRF4 rol-6(d))*

SM2544 *pxEx627 (let-413::GFP, pRF4 rol-6(d)); set-25(n5021) III* KM267 *HS::hlh-1*

SM2441 *HS::hlh-1; set-25(n5021) III*

SM2333 *pxSi01 (zen-4::gfp, unc-119+) II; unc-119(ed3) III*.

#### Immunofluorescence

Embryos were collected, fixed and stained according to Kiefer et al. 2007. <u>Primary antibodies</u>: H3K9me2 (1:200) Abcam ab1220; H3K9me3 (1:200) Kimura 2F3; H3K27me3 (1:200) Active Motif 61017; Pan-histone (1:500) Chemicon/Millipore MAB052; Histone H3 (1:500) Abcam ab1791; DLG-1 (1:200) Thermo Scientific PSD95; Paramyosin (1:50) Developmental Studies Hybridoma Bank 5–23 <u>Secondary antibodies</u>: Goat ±-mouse IgG Alexa 488 and 594 (1:200) (Abcam ab150117, ab150116); Goat ±-rabbit IgG Alexa 488 and 594 (1:200) (Abcam ab150077, ab150080)

<u>Quantitation of images</u>: To avoid slide to slide variation, wild-type and *set-25* mutant embryos were placed on the same slide. WT embryos carried a single copy of integrated ZEN-4::GFP (Von Stetina et al. 2017), to distinguish WT from *set-25* embryos.

<u>Image analysis</u>: Stacks of optical sections were collected with a ZEISS LSM700 Confocal Microscope, and analyzed using Volocity Software (PerkinElmer). Nuclei were identified in 3-D using DAPI, and signal intensity of histone modifications was calculated for each nucleus. Mitotic nuclei, based on morphology, were excluded from analysis. Mean cytoplasmic background was measured for each embryo and subtracted from the nuclear signal. For each nucleus, signal intensity of the histone modification was normalized to signal intensity of pan histone or total histone H3.

#### FAIRE-seq

FAIRE was adapted from (Giresi et. al, 2007). Briefly, 100,000 embryos were harvested and frozen. Frozen embryos were thawed on ice. Embryos underwent rapid freezing in liquid nitrogen followed by thawing to crack eggshell and achieve uniform crosslinking. To dislodge unstable protein-DNA interactions, embryos were incubated in 1X M9 (85mM NaCl, 42mM Na_2_HPO_4_, 22mM NaH_2_PO_4_, 2mM MgSO_4_, 1X EDTA-free proteinase inhibitor (Millipore 539134), 1X PhosStop (Roche)) at 4°C for 30min with rotation. Embryos were fixed in 1.5% formaldehyde in M9 buffer at 20°C for 30min with constant rotation and quenched with 125mM glycine. Embryos were washed 3x with ice cold M9 before adding lysis buffer (50mM Tris pH8.0, 10M EDTA pH8.0, 0.2% SDS) and incubated on ice for 10min. Embryos were sonicated with Q-sonica Q700 cup horn system at 80% amplitude, with cycles of 10 seconds on + 20 seconds off, resulting in fragment sizes between 100–500bp (average ∼200bp). 50ul of the lysate was used for Input DNA and 200ul for FAIRE DNA. FAIRE sample was treated with 5ul of 10mg/ml proteinase K at 60C for 16hr. Both input and FAIRE samples were subjected to phenol-chloroform extraction twice and chloroform once. Samples were ethanol precipitated and dissolved in 50ul TE buffer, and treated with RNase A. Samples were purified with Qiagen PCR clean up kits to remove short nucleotides prior to measuring the DNA concentration with Qubit DNA HS kit. FAIRE was performed 3 times for WT and *set-25* late-stage embryos. FAIRE recovery is sensitive to embryo growth, fixation and sonication Therefore, we preformed FAIRE on embryo preps that had good viability (>90% hatch rate after harvesting), and performed comparison experiments side-by-side and with C. briggsae normalization.

FAIRE DNA and Input DNA were size selected for 100–600bp, which included essentially all FAIRE DNA, to limit sampling bias. Sequencing libraries were generated from 10ng of DNA with the TruSeq kit (Illumina IP-202–1020) following manufactures’ guidelines. PCR amplification was limited to 10 cycles to minimize PCR-related bias. Libraries were sequenced with Illumina HiSeq 2000 using 50nt single-read setting.

Sequencing reads were mapped to the *C. elegans* reference genome version WS220 with BWA. Reads that mapped to multiple genomic regions with equal scores were randomly assigned to one region. Mapped reads were piled up using MACS2 callpeak after extending the single-end reads for 200bps towards its 3’ direction, and normalized by the sequencing depth to compare different samples. Sequence data is in process of being submitted to GEO.

#### 5 state HMM analysis

To identify regions with differential FAIRE signal between WT and *set-25* mutant embryos, 5-state Hidden Markov model implemented in R was applied. The BWA mapped bam files were converted to coverage bedgraphs using bedtools genomecov with a 200 bp fragment size. These coverage bedgraphs were the input files. The HMM model uses a binomial probability model where the number of reads from the *set-25* mutant embryos were treated as the number of successes, and the probability of success (i.e, the probability of picking a mutant read from the total reads mapping to that position in the genome) varies across states. This was effectively equivalent to the proportion of reads covering each base that were from the *set-25* mutant sample relative to the WT. Coverage of every 10 base pairs were sampled across the genome and then we separately estimate the transition probabilities and the binomial probabilities for each of the five states for each chromosome using the Baum Welch algorithm. We then infer which state each position belongs to using the Viterbi algorithm, both from the Hidden Markov package. Finally, we covert these state descriptions into bed files using a custom Python script that gets the start and end position of contiguous runs of 1 or more positions in the same state. These bed files are the main input to subsequent analysis.

Many of the initial state 5 regions were separated by small state 4 regions. To reduce the complexity, these adjacent state 5 regions were merged with bedtools merge, and retained the intervals that contained regions that were entirely state 5, or merged state 4 and state 5. This final state 5 file was used for downstream analysis.

#### High-density map for repeats and H3K9 methylated regions

Repeat annotation was obtained from RepeatMasker on UCSC table browser, WS220 genome version. This file was separated to repeat classes; DNA transposons, RNA transposons and satellite repeats, based on the repClass. H3K9me2 (GSE49736) and H3K9me3 (GSE49732) methylated regions were defined by MACS2 peak call with parameters ‘-g ce -q 0.05 -extsize 146’. High-density scatter maps were created with R, ggplots2 package.

#### Heat map and cell type enriched TSS

Cell-type enriched genes were defined by Spencer (Spencer, Mill, 2011, Genome Research). TSS files used in the heat maps were the most confident TSSs, that were reported by both Chen and Krusei. Heat maps were generated with R, heatmap2 package. For each gene, the average FAIRE value were calculated for 50bp per window, 40 windows up stream and 40 windows downstream. K-mean cluster and heat maps were generated with Cluster and TreeView program, respectively.

#### Single molecule fluorescence in situ hybridization for *dlg-1* mRNA

smFISH procedure was adopted from (Raj et al. 2008) with modifications. *dlg-1, CeRep55* and *Mirage-1* smFISH probes labeled with Qusar570 were designed by the Stellaris FISH Probe Designer (Biosearch Technologies. Sequences available upon request). Probes were dissolved in 400ml TE (10mM Tris pH 8.0, 1mM EDTA pH 8.0) to 0.2 nmole. Embryos were dissected from gravid worms in embryo buffer, and incubated at 20°C for 4hr to reach late-stage before transferred to poly-L-lysine (Sigma P-8920) coated glass slides. Embryos were then fixed in 100ul 3.7% formaldehyde in 1X PBS for 15 minutes at RT, followed by the freeze crack method to break the egg shell (Duerr 2006).The slides were immersed in 70% EtOH and stored at 4°C for 16hr. Slides were washed with wash buffer (10% formamide in 2X SSC). 100ul of hybridization buffer (10% formamide and 10% dextran sulfate in 2X SSC) containing 0.001 nmol smFISH probe was added to the embryos, and incubated for 4 hours at 37°C in a humidifier chamber. The embryos were rinsed once quickly using wash buffer, then washed for 1h at 37°C. 7ul of SlowFade with DAPI (Thermo Fisher) was added to the slides and sealed with nail polish. Images were obtained with Zeiss fluorescence microscope with ApoTome attachment on 63X oil objectives. Selected single plane images that encompassed the connected arcade cells from ZEN Blue image software (Zeiss) were imported into ImageJ for quantification. Let-413::GFP signal was used to trace the cell boundary of arcade cells and the arcade lumen. The background noise of *dlg-*1 smFISH signal was removed by rolling ball method with 100 pixel radius.

#### Viability assays

**Embryo hatching rate**. Embryos were dissected from gravid adults in 1X M9. 25 embryos were transferred to a 3.5cm diameter plate with a small OP50 bacterial lawn in the center. 4 plates per genotype per experiment and repeated three times. Hatching rate was recorded 24hr after embryos were transferred. **Embryo development time**. Embryos were dissected from gravid adults in 1X M9, and 2-cell stage embryos were transferred to a 3.5cm diameter plate without food. After 10hr plates were observed under dissection microscope every 10min, until the last embryo hatched. **Brood size**. L4 worm was singled onto a 6cm diameter plate with OP50 bacterial lawn. Embryos that were laid on the plates were counted and the original worm was transferred to a new plate. Repeated counting and transferring for 3–4 days, when the adult worm stopped producing embryos. Life span. L4 worm was singled onto a 6cm diameter plate with OP50 bacterial lawn, and was transferred to a new plate after 3 days and again after another 3 days. When the worm stopped moving, i.e. no crawling and no pharynx pumping, it was pronounced dead. Animals used in the above viability assays were out-crossed to wild-type or *set-25* and siblings used as controls.

#### Hlh-1 challenge assay

Embryos were dissected from gravid adults (KM267, SM2441) in embryo buffer. 2-cell stage embryos were transferred to poly-L-lysine coated slide and incubated in a humidify chamber at 22.5°C for 4.5–5hr, when embryos reached comma-stage. Embryos that were younger or older than comma stage were removed from the slides. The comma-stage embryos were transfer to a pre-heated 34°C humidify chamber for 10min, and rapidly returned to a 20°C humidify chamber to resume embryogenesis for 5hr (intermediate phenotype) or for 16hr (terminal phenotype).

#### Published data used in this paper

Pol II: GSM1666982; H4K8ac: GSE49737; H3K79me3: GSE49743; H3K79me2: GSE49742; H3K79me1: GSE49741; H3K36me3: GSE50264; H3K36me1: GSE49740; H3K27ac: GSE49734; H3K4me3: GSE49739; H3K4me2: GSE49733; H3K4me1: GSE50262; H3K27me3: GSE49738; H3K4me3: GSE49739; H3K9me1: GSE49744; H3K9me2: GSE49736; H3K9me3: GSE49732

Epithelia and Muscle gene sets are from (Spencer et al. 2011).

## Acknowledgments

We thank J. Gaspar and C. Aydin for bioinformatics, A. Schier and R. Losick for comments on the manuscript, and the Lieb lab for input on FAIRE. BM was supported by AAUW. BM, SEM and HMC were supported by 5R37GM056264 and Harvard Univ.

## Author Contributions

HMC developed FAIRE and performed the profiling and heat-map analysis, and wrote some of the text, TBS developed HMM; TBS, TL, EL and JW performed computational analysis. BM performed heat-shock experiments and contributed to writing, SKR performed smFISH; SEM contributed financial support and intellectual guidance, and wrote portions of the paper.

